# Inhibition of the JAK/STAT pathway with baricitinib reduces the multiple organ dysfunction caused by hemorrhagic shock in rats

**DOI:** 10.1101/2022.01.13.476088

**Authors:** Nikita Mayur Patel, Debora Collotta, Eleonora Aimaretti, Gustavo Ferreira Alves, Sarah Kröller, Sina Maren Coldewey, Massimo Collino, Christoph Thiemermann

## Abstract

**Objective:** The aim of this study was to investigate (a) the effects of the Janus kinase (JAK)/signal transducer and activator of transcription (STAT) pathway inhibitor (baricitinib) on the multiple organ dysfunction syndrome (MODS) in a rat model of hemorrhagic shock (HS) and (b) whether treatment with baricitinib attenuates the activation of JAK/STAT, NF-κB and NLRP3 caused by HS.

**Background:** Post-traumatic MODS, which is in part due to excessive systemic inflammation, is associated with high morbidity and mortality. The JAK/STAT pathway is a regulator of numerous growth factor and cytokine receptors and, hence, is considered a potential master regulator of many inflammatory signaling processes. However, its role in trauma-hemorrhage is unknown.

**Methods:** An acute HS rat model was performed to determine the effect of baricitinib on MODS. The activation of JAK/STAT, NF-κB and NLRP3 pathways were analyzed by western blotting in the kidney and liver.

**Results:** We demonstrate here for the first time that treatment with baricitinib (during resuscitation following severe hemorrhage) attenuates the organ injury and dysfunction and the activation of JAK/STAT, NF-κB and NLRP3 pathways caused by HS in the rat.

**Conclusions:** Our results point to a role of the JAK/STAT pathway in the pathophysiology of the organ injury and dysfunction caused by trauma/hemorrhage and indicate that JAK inhibitors, such as baricitinib, may be repurposed for the treatment of the MODS after trauma and/or hemorrhage.

## Introduction

Trauma is one of the major causes of mortality in those aged under 44 and the number of patients dying from trauma surpasses those from tuberculosis, malaria and HIV/AIDS combined^1^. Annually, there are approximately 6 million trauma-associated mortalities worldwide. Trauma-associated hemorrhage accounts for almost 40% of all trauma deaths^2^ and is a significant driver of multiple organ dysfunction syndrome (MODS)^3^. There is good evidence that a) uncontrolled systemic inflammation secondary to the release of damage-associated molecular patterns (DAMPs) from substantial tissue damage and b) ischemiareperfusion injury contribute to the onset of MODS^4^. However, there are no specific pharmacological interventions clinically used to prevent the onset of HS-induced MODS.

Circulating pro-inflammatory mediators bind to cell-surface receptors and subsequently signal through activation of intracellular protein tyrosine kinases, including the Janus kinase (JAK) family. The JAK family is composed of JAK1, JAK2, JAK3 and tyrosine kinase 2 (TYK2) and these isoforms interact with the signal transducer and activator of transcription (STAT) class of proteins^5^. JAK inhibitors affect the JAK-STAT protein interaction and disrupt the propagation of signals through the JAK/STAT pathway, thus modulating inflammatory signaling processes.

We have recently shown that JAK/STAT inhibition reduced both the diet-related metabolic derangements and the associated organ injury/dysfunction in a murine model of high-fat diet-induced type 2 diabetes^6^. Specifically, treatment with baricitinib, a JAK1/JAK2 inhibitor, resulted in improvements of diet-induced myosteatosis, mesangial expansion and associated proteinuria in addition to reducing blood cytokine levels, renal dysfunction and skeletal muscle damage.

Driven by the COVID-19 pandemic, strategies focusing on drug repurposing which dampen the cytokine storm and pulmonary injury associated with severe acute respiratory syndrome coronavirus 2 (SARS-CoV-2) have been examined. Recent investigations have shown that baricitinib, which is thought to possess antiviral properties through its affinity for adaptor-associated kinase-1 and the subsequent reduction in SARS-CoV-2 endocytosis, improves clinical status in patients with COVID-19^7–21^. Thus, baricitinib appears to lower systemic inflammation and several ongoing clinical trials are investigating the potential impact of this repurposing approach on outcome in COVID-19 patients (ClinicalTrials.gov Identifier: NCT04320277, NCT04321993, NCT04346147, NCT04381936, NCT04390464, NCT04399798, NCT04640168, NCT04693026, NCT04832880, NCT04890626, NCT04891133, NCT04970719, NCT05056558, NCT05082714, NCT05074420, NCT04393051).

Baricitinib is used in patients for the treatment of moderate-to-severe active rheumatoid arthritis and atopic eczema/dermatitis, with the latter use approved by the NICE in 2021^22^. Given the evident beneficial effects of baricitinib administration in type 2 diabetes and COVID-19, we wished to explore the potential of repurposing baricitinib in trauma-hemorrhage. Currently, there is limited information about the role of JAK/STAT pathway in trauma^23–25^.

## Methods

### JAK2 and STAT3 gene expression in human whole blood

Original data were obtained under Gene Expression Omnibus (GEO) accession GSE36809, published by Xiao and colleagues^26^. RNA was extracted from whole blood leukocytes of severe blunt trauma patients (n = 167) over the course of 28 days and healthy controls (n = 37) and hybridized onto an HU133 Plus 2.0 GeneChip (Affymetrix) according to the manufacturer’s recommendations. The dataset was reanalyzed for JAK2 and STAT3 gene expression. A further reanalysis was performed dividing the trauma patients into uncomplicated (n = 55) and complicated (n = 41) groups.

### Use of Experimental Animals - Ethics Statement

All animal procedures were approved by the Animal Welfare Ethics Review Board of Queen Mary University of London and by the Home Office (License number PC5F29685).

### Experimental Design

Male Wistar rats (Charles River Laboratories Ltd., Kent, UK) weighing 260-310 g were kept under standard laboratory conditions and received a chow diet and water *ad libitum*. Baricitinib (Insight Biotechnology, UK) was diluted in 5 % DMSO + 95 % Ringer’s Lactate (vehicle) and rats were treated (i.p.) upon resuscitation. Further information about baricitinib can be found in the supplemental.

### Hemorrhagic Shock Model

The acute hemorrhagic shock model was performed as previously described^27,28^. Briefly, forty rats were anesthetized with sodium thiopentone (120 mg/kg i.p. initially and 10 mg/kg i.v. for maintenance as needed) and randomized into four groups: Sham + vehicle (n = 10); Sham + baricitinib (1 mg/kg; n = 9), HS + vehicle (n = 11); HS + baricitinib (1 mg/kg; n = 10). Blood was withdrawn to achieve a fall in mean arterial pressure (MAP) to 35 ± 5 mmHg, which was maintained for 90 min. At 90 min after initiation of hemorrhage (or when 25% of the shed blood had to be reinjected to sustain MAP), resuscitation was performed. At 4 h post-resuscitation, blood was collected for the measurement of biomarkers of organ injury/dysfunction (MRC Harwell Institute, Oxfordshire, UK). Sham-operated rats were used as control and underwent identical surgical procedures, but without hemorrhage or resuscitation. Detailed description of the model can be found in the supplemental (Supplementary eFigure 1).

### Western Blot Analysis

Semi-quantitative immunoblot analysis was carried out in liver and kidney tissue samples as previously described^27,28^. Detailed description of the method can be found in the supplemental.

### CD68 immunohistochemical staining

Renal sections (2 μm) were deparaffinized and hydrated as previously described^29^ and stained for CD68. Detailed description of the method can be found in the supplemental.

### Periodic acid Schiff staining

Periodic acid Schiff (PAS) staining of renal sections (2 μm) was performed using a PAS staining kit (Carl Roth, Karlsruhe, Germany). Histomorphological changes were evaluated at 20x magnification using a scoring system as previously described^29^. Images were taken using a KEYENCE BZ-X800 microscope and BZ-X800 viewer after performing white balance and auto exposure.

### Statistical Analysis

All data in text and figures are expressed as mean ± SEM of *n* observations, where *n* represents the number of animals/experiments/subjects studied. Measurements obtained from the patient groups and vehicle and baricitinib treated animal groups were analyzed by one-way/two-way ANOVA followed by Bonferroni’s *post-hoc* test or Kruskal-Wallis test followed by Dunn’s *post-hoc* test on GraphPad Prism 8.0 (GraphPad Software, Inc., La Jolla, CA, USA). The distribution of the data was verified by Shapiro-Wilk normality test, and the homogeneity of variances by Bartlett test. When necessary, values were transformed into logarithmic values to achieve normality and homogeneity of variances. P<0.05 was considered statistically significant.

## Results

### JAK2 and STAT3 gene expression is elevated in trauma patients

Xiao and colleagues^26^ compared genome-wide gene expression in leukocytes from trauma patients against matched healthy controls. We reanalyzed this dataset for JAK2 and STAT3 expression, as STAT3 is the most commonly stimulated downstream STAT following JAK2 activation. When compared to healthy controls, JAK2 expression was significantly elevated in patients with trauma at all time points (p<0.05; Figure 1A). An initial peak was noted at Day 1 followed by a gradual decrease, however, JAK2 expression still remained elevated at Day 28 after trauma. Similarly, STAT3 expression was significantly increased at all time points (p<0.05; Figure 1B) and an initial peak was noted at 12 h followed by a gradual decrease. However, STAT3 expression also still remained elevated at Day 28 after trauma. When comparing trauma patients stratified into uncomplicated (recovery in <5 days) and complicated (recovery after 14 days, no recovery by Day 28 or death) groups, JAK2 expression was significantly higher on Days 4 and 7 (p<0.05; Figure 1C) and STAT3 expression was significantly raised at 12 h (p<0.05; Figure 1D) in complicated patients when compared to uncomplicated trauma patients. These data indicate that the JAK2/STAT3 pathway may play a role in the pathophysiology of trauma.

**Figure 1:**
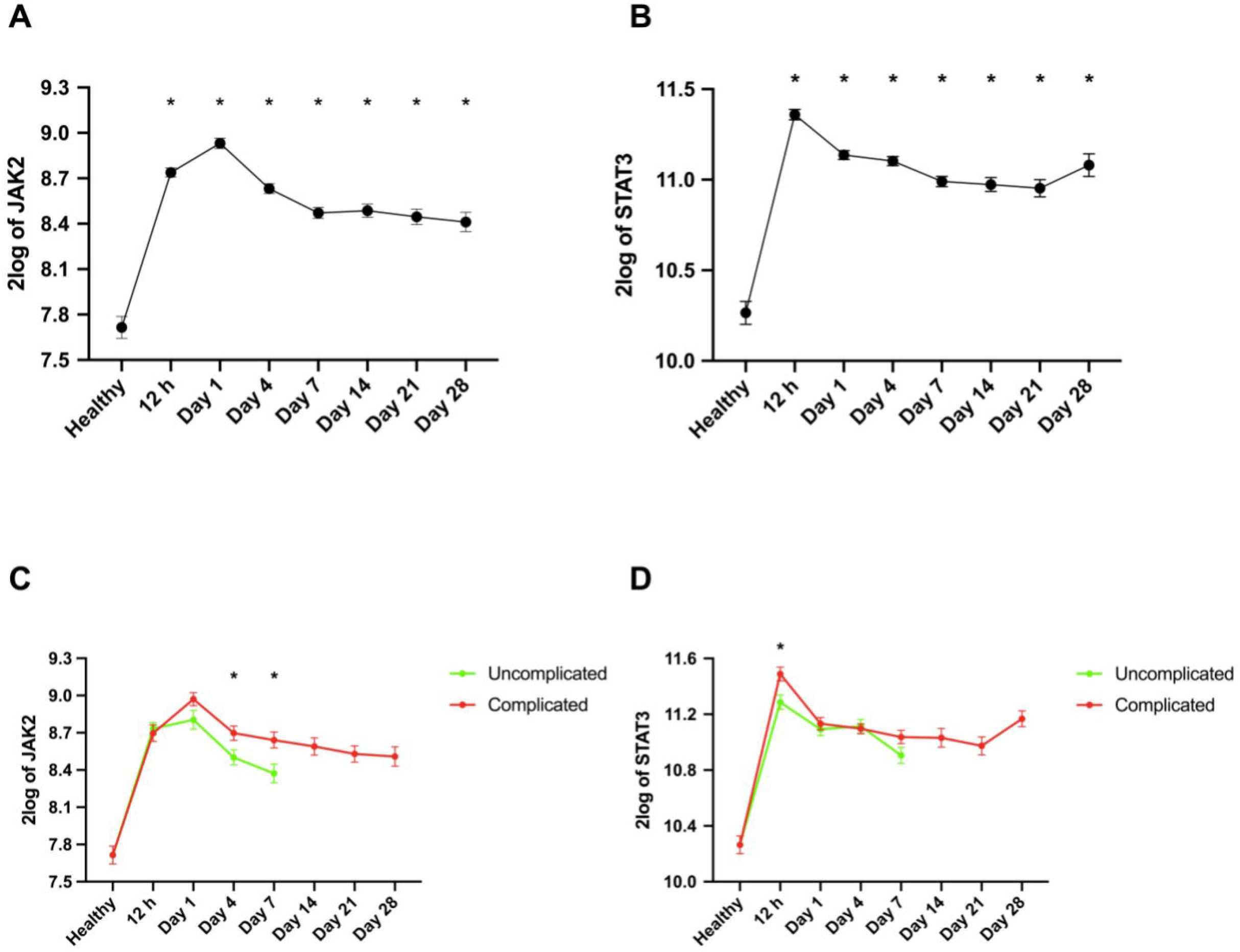
JAK2 and STAT3 gene expression is elevated in trauma patients. Original data was obtained from the Gene Expression Omnibus under dataset accession number GSE36809 which was published by Xiao and colleagues ^26^. RNA was extracted from whole blood leukocytes over a 28-day time course from trauma patients (n = 167) and matched healthy controls (n = 37). Data were reanalyzed for (**A**) JAK2 and (**B**) STAT3 gene expression in trauma patients and (**C**, **D**) uncomplicated vs. complicated trauma patient groups. Data are expressed as mean ± SEM. Statistical analysis was performed using one-way or two-way ANOVA followed by a Bonferroni’s *post-hoc* test. *p<0.05 denoted statistical significance.

### Baricitinib attenuates HS-induced organ injury/dysfunction in HS

Here we explored whether pharmacological intervention with the JAK1/JAK2 inhibitor baricitinib attenuates the MODS associated with HS in rats. When compared to sham-operated rats, rats subjected to HS displayed significant increases in serum urea (p<0.05; Figure 2A) and creatinine (p<0.05; Figure 2B); indicating the development of renal dysfunction. When compared to sham-operated rats, vehicle treated HS-rats exhibited significant increases in ALT (p<0.05; Figure 2C) and AST (p<0.05; Figure 2D) indicating the development of hepatic injury, while the increases in amylase (p<0.05; Figure 2E) and CK (p<0.05; Figure 2F) denote pancreatic and neuromuscular injury, respectively. The significant increase in LDH (p<0.05; Figure 2F) in vehicle treated HS-rats confirmed tissue injury. Treatment of HS-rats with baricitinib significantly attenuated the renal dysfunction, hepatic injury, pancreatic injury, neuromuscular injury and general tissue damage caused by HS as shown by the reduction in serum parameter values (all p<0.05; Figures 2A-G). Administration of baricitinib to sham-operated rats had no significant effect on any of the parameters measured (p>0.05; Figure 2).

**Figure 2:**
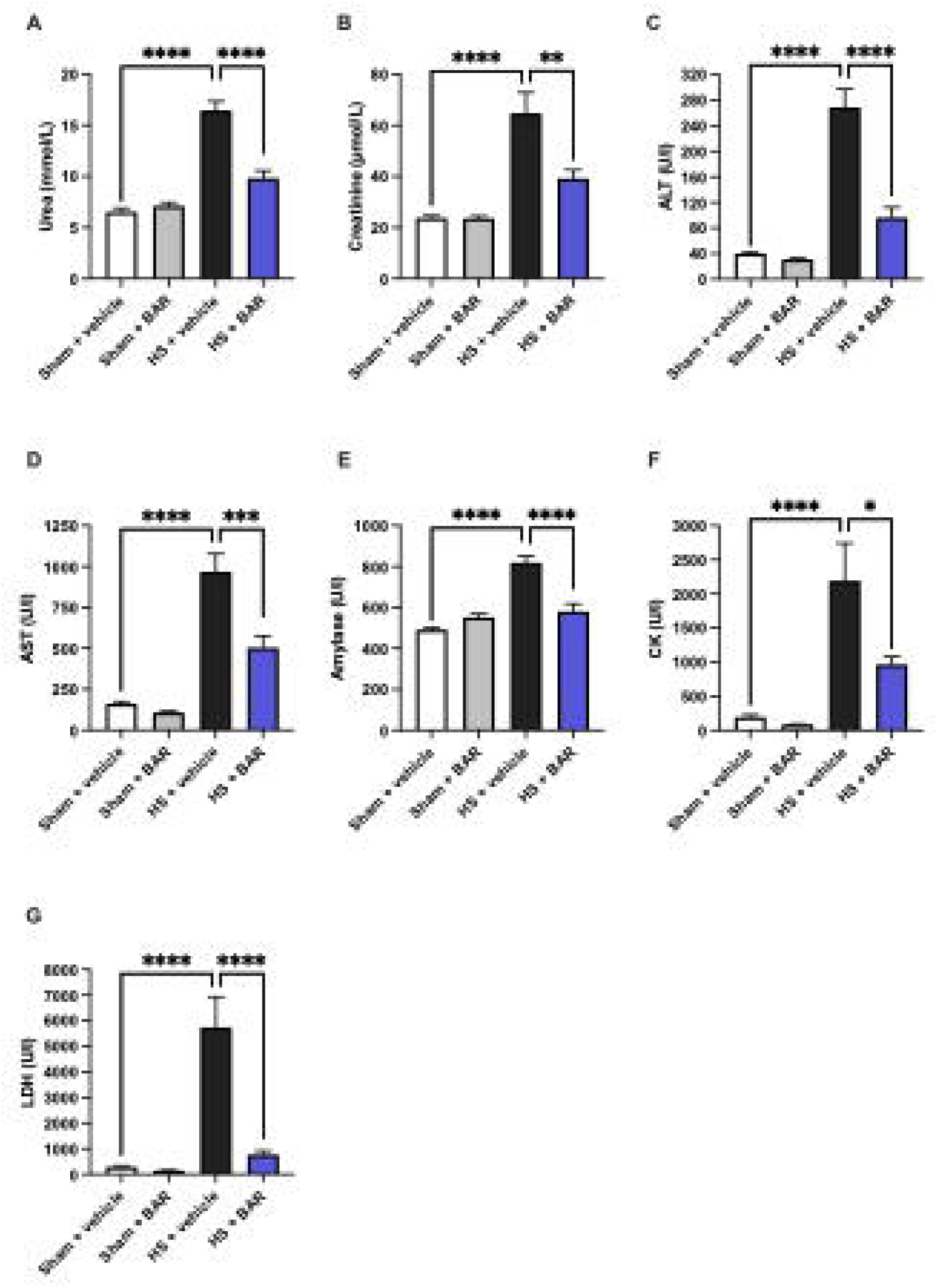
Baricitinib attenuates HS-induced organ injury/dysfunction in HS. Rats were subjected to hemorrhagic shock (HS) and 4 h after resuscitation, levels of serum (**A**) urea, (**B**) creatinine, (**C**) alanine aminotransferase (ALT), (**D**) aspartate aminotransferase (AST), (**E**) amylase, (**F**) creatine kinase (CK) and (**G**) lactate dehydrogenase (LDH) were determined in vehicle and BAR (baricitinib) treated rats. Sham-operated rats were used as control. Data are expressed as mean ± SEM of 9-11 animals per group. Statistical analysis was performed using one-way ANOVA followed by a Bonferroni’s *post-hoc* test. *p<0.05 denoted statistical significance.

### Baricitinib abolishes hepatic and renal JAK/STAT activation in HS

Using western blot analysis, we examined whether HS leads to the activation of the JAK2/STAT3 pathway in the liver and kidney, given that treatment with baricitinib significantly attenuated HS-associated hepatic injury and renal dysfunction. When compared to sham-operated rats, vehicle treated HS-rats displayed significant increases in the phosphorylation of hepatic and renal Jak2 at Tyr^1007-1008^ (p<0.05; Figures 3A-B) and Stat3 at Tyr^705^ (p<0.05; Figures 3C-D), indicating that the JAK2/STAT3 pathway is activated in the injured liver and kidneys. Treatment of HS-rats with baricitinib significantly abolished these increases in JAK2/STAT3 phosphorylation (p<0.05; Figures 3A-D).

**Figure 3:**
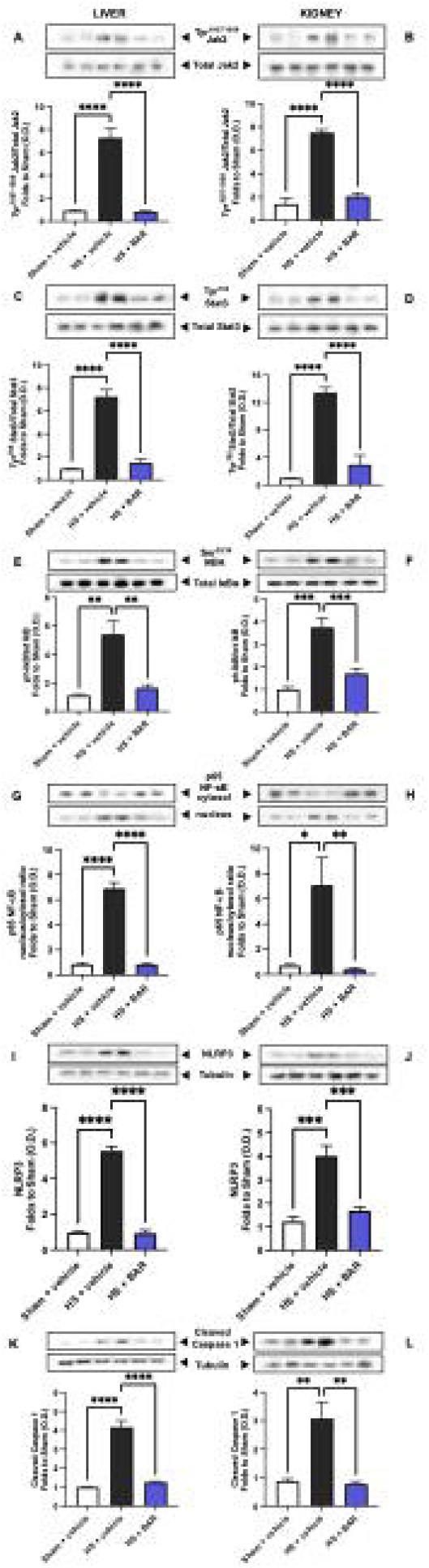
Baricitinib abolishes JAK/STAT, NF-κB and NLRP3 activation in HS. (**A-B**) The phosphorylation of Jak2 at Tyr^1007-1008^, (**C-D**) the phosphorylation of Stat3 at Tyr^705^, (**E-F**) the phosphorylation of IκBα at Ser^32/36^, (**G-H**) the nuclear translocation of p65, (**I-J**) the activation of NLRP3 and (**K-L**) the cleaved (activated) form of caspase 1 of vehicle and BAR (baricitinib) treated rats were determined by western blotting in the liver and kidney. Protein expression was measured as relative optical density (O.D.) and normalized to the sham band. Data are expressed as mean ± SEM of 4-5 animals per group. Statistical analysis was performed using one-way ANOVA followed by a Bonferroni’s *post-hoc* test. *p<0.05 denoted statistical significance.

### Baricitinib abolishes hepatic and renal NF-κB activation in HS

The effect of JAK/STAT inhibition on the activation of NF-κB were investigated in liver and kidney. When compared to sham-operated rats, vehicle treated HS-rats had significant increases in the hepatic and renal phosphorylation of IκBα at Ser^32/36^ (p<0.05; Figures 3E-F) and the translocation of p65 to the nucleus (p<0.05; Figures 3G-H). Treatment of HS-rats with baricitinib significantly abolished this activation of NF-κB (p<0.05; Figures 3E-H).

### Baricitinib abolishes hepatic and renal NLRP3 and caspase 1 activation in HS

Having discovered that baricitinib significantly reduced NF-κB activation in the liver and kidney of HS-rats, we next analyzed the potential involvement of the NLRP3 inflammasome complex. When compared to sham-operated rats, vehicle treated HS-rats exhibited significantly increased hepatic and renal expression of the NLRP3 inflammasome (p<0.05; Figures 3I-J) and cleaved (activated) form of caspase 1 (p<0.05; Figures 3K-L). Treatment of HS-rats with baricitinib significantly inhibited these increases (p<0.05; Figures 3I-L).

### JAK2 activation correlates with hepatic injury, renal dysfunction, STAT3, NF-κB and NLRP3 activation in an acute HS model

Correlation analysis was performed to determine whether the degree of activation of JAK correlates with changes in liver condition, renal function, STAT3, NF-κB and NLRP3 activation. Significant positive correlations were found between JAK2 activation and (with the exception of AST) all parameters investigated (Supplementary eFigure 2).

### Baricitinib reduces renal macrophage invasion and tissue injury in HS

Having shown that baricitinib significantly attenuated the HS-induced renal dysfunction, we investigated renal macrophage infiltration and tissue injury determined by CD68 and PAS staining, respectively. When compared to sham-operated rats, vehicle treated HS-rats exhibited a significantly increased expression of CD68 in the kidney (p<0.05; Figure 4A). Treatment with baricitinib in HS-rats significantly reduced CD68 expression, indicating an attenuation of renal macrophage invasion (p<0.05; Figure 4A). When compared to sham-operated rats, vehicle treated HS-rats displayed a significantly raised renal PAS score (p<0.05; Figure 4C). Treatment with baricitinib in HS-rats reduced this rise in PAS score (albeit this change was not statistically significant), suggesting a tissue protective effect of baricitinib in the kidney (Figure 4C).

**Figure 4:**
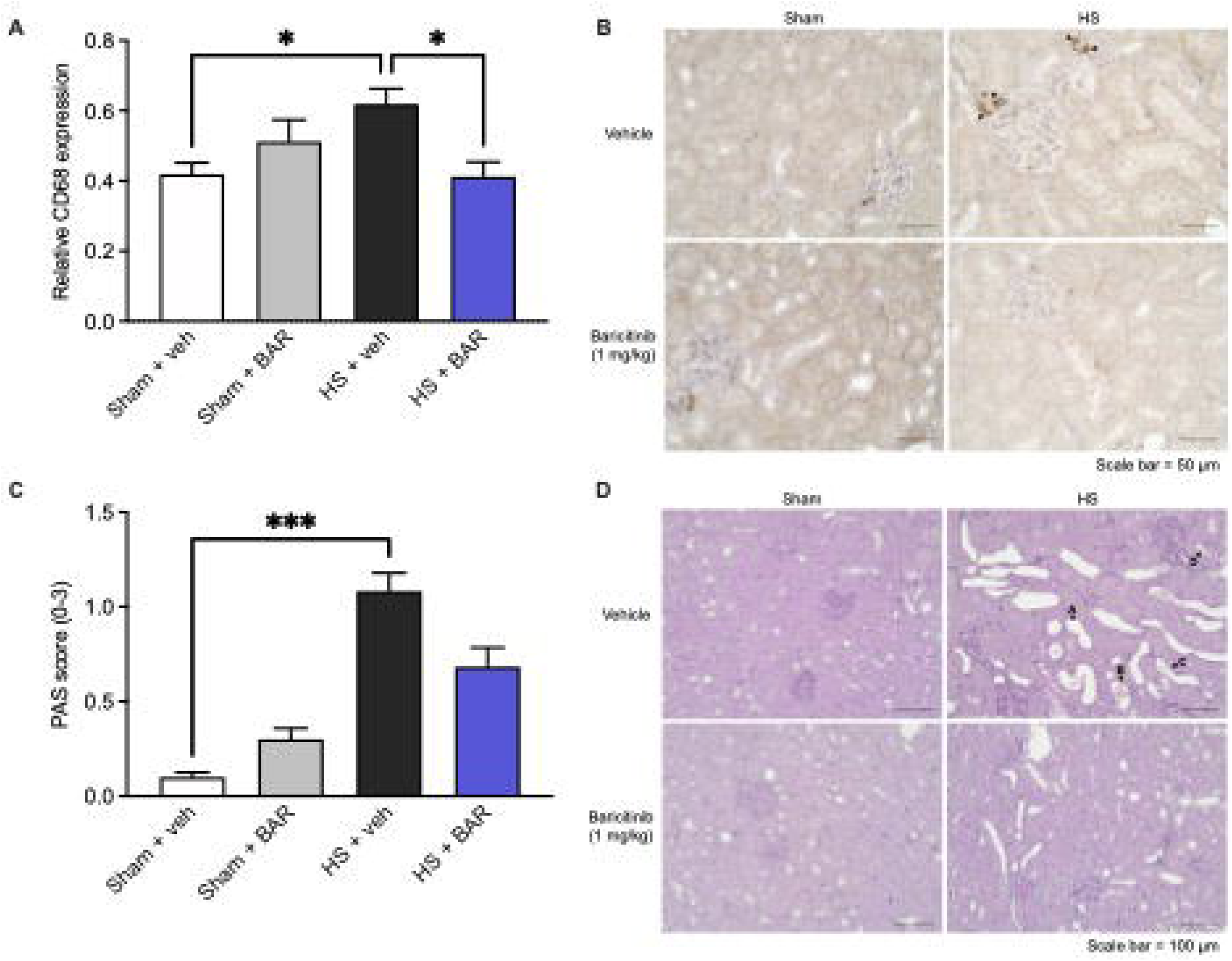
Baricitinib reduces renal macrophage invasion and tissue injury in HS. Rats were subjected to hemorrhagic shock (HS) and 4 h after resuscitation, (**A**) relative CD68 expression and (**C**) Periodic acid Schiff (PAS) scores (Score 0: no damage, 1: <25% damaged, 2: 25-50% damaged and 3: >50% damaged) were quantified in renal tissue of vehicle and BAR (baricitinib) treated rats. Representative images of (**B**) CD68 staining (40x magnification) and (**D**) PAS staining (20x magnification) are shown. Black arrows highlight (**B**) sites of positive CD68 staining and (**D**) tubule dilatation (arrow A), detached cell material (arrow B), flattened epithelial cells (arrow C) and beginning stages of tubular brush border loss in proximal tubules (arrow D). Data are expressed as mean ± SEM of eight animals per group. Statistical analysis was performed using (**A**) one-way ANOVA followed by a Bonferroni’s *post-hoc* test and (**C**) Kruskal-Wallis test followed by a Dunn’s *post-hoc* test. *p<0.05 denoted statistical significance.

### Effect of baricitinib on HS-induced circulatory failure in HS

To investigate the effects of baricitinib on circulatory failure, MAP was measured from the completion of surgery to the termination of the experiment. Baseline MAP values were similar amongst all four groups. Rats subjected to HS demonstrated a decline in MAP which was ameliorated by resuscitation, but MAP remained lower than that of sham-operated rats during resuscitation (at the equivalent time points, Figure 5). The MAP of baricitinib treated HS-rats was similar to those of vehicle treated HS-rats at the end of the resuscitation period (Figure 5). Administration of baricitinib to sham-operated rats had no significant effect on MAP (p>0.05; Figure 5).

**Figure 5:**
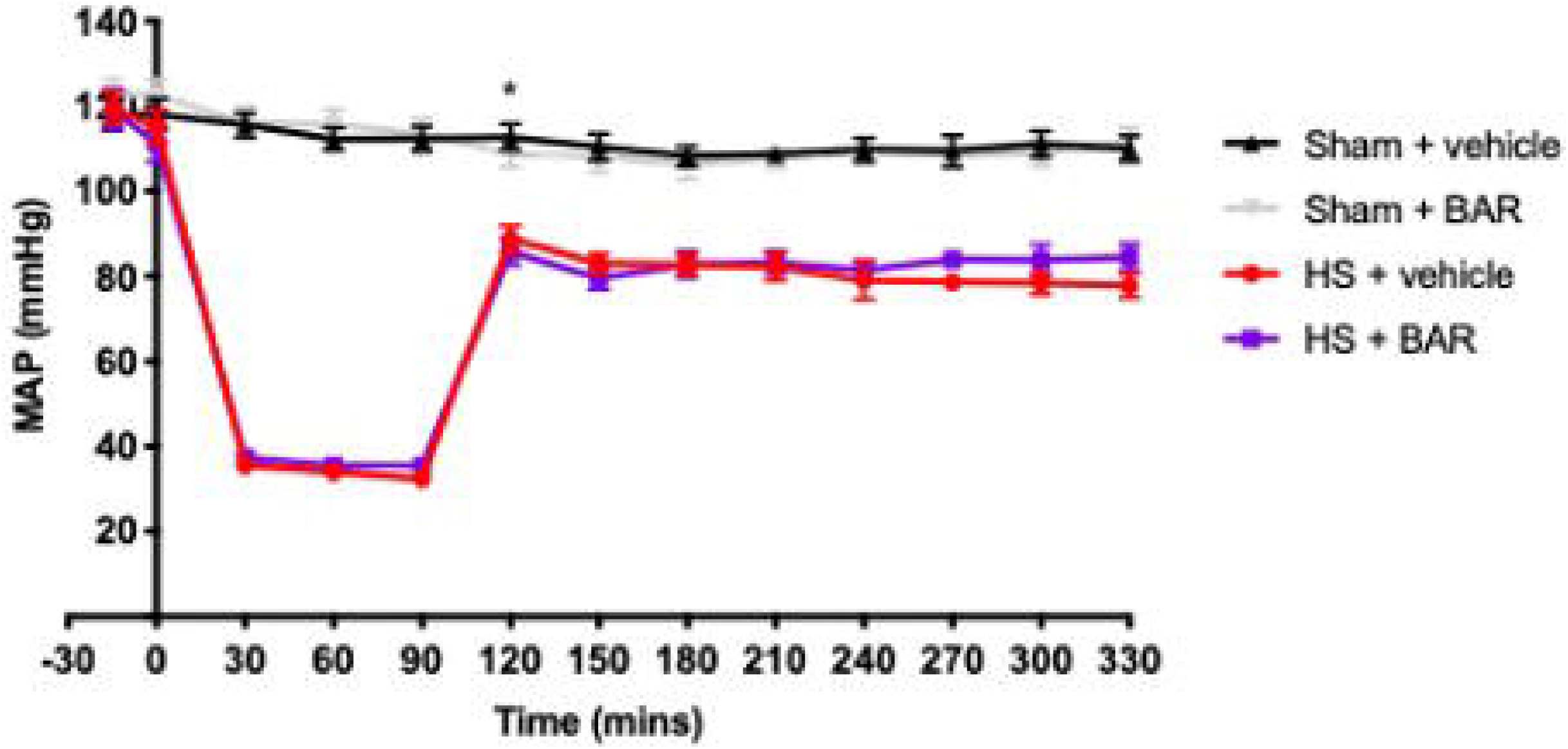
Effect of baricitinib on HS-induced circulatory failure in HS. Mean arterial pressure (MAP) was measured from the completion of surgery to the termination of the experiment for vehicle and BAR (baricitinib) treated rats. Data are expressed as mean ± SEM of 9-11 animals per group. Statistical analysis was performed using two-way ANOVA followed by a Bonferroni’s *post-hoc* test. *p<0.05 Sham + vehicle vs. HS + vehicle.

## Discussion

This study reports that inhibition of JAK/STAT activity attenuates the organ injury/dysfunction associated with HS (acute rat model; Figure 2). Having shown that JAK2 and STAT3 gene expression is significantly elevated in leukocytes of trauma patients (Figures 1A-B) and, most notably, that the expression of JAK2 and STAT3 is higher in trauma patients with a complicated recovery (Figures 1C-D), we used a reverse translational approach to investigate whether pharmacological intervention with the JAK1/JAK2 inhibitor baricitinib ameliorates the MODS associated with HS in a well-established rat model^30–32^.

Baricitinib inhibits JAK1 and JAK2 to an equal degree and is currently used in patients with inflammatory diseases, including rheumatoid arthritis and atopic dermatitis^33^. We report here for the first time that baricitinib significantly attenuated the renal dysfunction, hepatic injury, pancreatic injury and neuromuscular injury caused by HS (Figure 2). Similarly, JAK inhibition lowers disease severity in animal models of ischemia-reperfusion injury^34–39^, sepsis^40–43^, acute kidney injury^44^ and endotoxemia^45^.

What, then, are the mechanisms by which baricitinib attenuates HS-associated organ injury/dysfunction? HS resulted in a significant increase in JAK2/STAT3 activity in the liver and kidneys (Figure 3) which correlated with the rises in ALT, creatinine and urea (Supplementary eFigure 2). Indeed, inhibition of JAK2/STAT3 activity with baricitinib in the liver and kidneys of HS-rats decreased the hepatic injury and renal dysfunction, suggesting that activation of the JAK2/STAT3 pathway plays a pivotal role in the pathophysiology of the MODS associated with HS. Despite not finding a significant correlation between JAK activation and AST in our study, genome-wide association study-prioritized genes and gene coexpression pattern analysis revealed JAK/STAT signaling was linked to AST-specific gene sets which supports the role of JAK/STAT in liver function^46^.

A synergistic interaction between STAT and NF-κB occur on many levels: these include physical interactions of the two molecules, cooperative binding at particular gene promoter subsets to induce target gene expression and cytokine feedback loops, whereby cytokines induced by STAT and NF-κB (e.g. IL-6) can further prolong the activation of both molecules^47^. Trauma leads to increased nuclear translocation of NF-κB^27,28,30–32^. Inhibition of JAK2/STAT3 activity with baricitinib decreased NF-κB activation in both the liver and kidneys (Figure 3). We also found a significant positive correlation between the activation of JAK2 and the phosphorylation of IκBα at Ser^32/36^ and translocation of p65 (Supplementary eFigure 2). This may suggest that blocking the activation of NF-κB contributes to the observed effects of baricitinib in HS. Activation of NF-κB drives the production of multiple pro- and anti-inflammatory mediators including enzymes, cytokines and chemokines^48^. As part of a positive feedback loop, these mediators can activate NF-κB and its upstream signaling components, further amplifying and sustaining the inflammatory responses mediated by NF-κB which can lead to greater permeability of the endothelium, hypoperfused/hypoxic tissues, tissue injury and eventually MODS^49^.

The JAK/STAT pathway has also been shown to be linked to the NLRP3 inflammasome, with pharmacological inhibition of JAK reducing the expression and activation of NLRP3 inflammasome components^50,51^. NLRP3 inflammasome activation drives the formation of IL-1β which plays a key role in the systemic inflammation and/or organ dysfunction associated with trauma^27,28^. Inhibition of JAK2/STAT3 activity with baricitinib decreased the assembly and successive activation of the NLRP3 inflammasome in the liver and kidneys (Figure 3). We also discovered a significant positive correlation between JAK2 activation and the activation of NLRP3 and caspase 1 (Supplementary eFigure 2). A recent phenotypic high-content, high-throughput screen identified targeting JAK to inhibit NLRP3 inflammasome activation^52^. This may suggest that blocking the activation of the NLRP3 inflammasome contributes to the observed protective effects of baricitinib in HS by reducing the pro-inflammatory effects of IL-1β and ensuing tissue inflammation.

The sterile inflammation caused by HS drives the recruitment of leukocytes to the tissues and is secondary to NF-κB and NLRP3 activation and their associated transcriptional regulation of pro-inflammatory cytokines^53–55^. Furthermore, leukocytes and endothelial cells express adhesion molecules, and this is regulated by NF-κB to facilitate leukocyte extravasation from the circulation to the site of damage^56^. It has been shown that experimental injury induces a transcriptomic shift in leukocytes 4 h post trauma and HS and several pathways related to innate immunity were upregulated; including those involved in myeloid leukocyte activation and differentiation^57^. We measured CD68 expression and PAS score in the kidney as markers of macrophage infiltration and overall tissue injury, respectively^58^. HS resulted in a significant rise in relative CD68 expression and PAS score in the kidney which was attenuated by baricitinib treatment in HS-rats (Figure 4, see also Supplement for an extended discussion). Please refer to the Supplement for a discussion of the study limitations.

Our results and conclusions are supported by findings in COVID-19 patient cohorts where the JAK/STAT pathway has been proposed to play a role in disease pathogenesis. The beneficial effects of baricitinib in COVID-19 patients, as measured by the improved levels of oxygenation, decreased need for mechanical ventilation (invasive and non-invasive), reduced ICU admission and mortality indicate that JAK/STAT activation contributes to the disease pathology^59^. Parallels can be drawn between COVID-19 and trauma-associated MODS, with both displaying features of an excessive systemic inflammatory state.

## Conclusions

In conclusion, we report here for the first time that baricitinib reduces the organ injury/dysfunction caused by severe hemorrhage in the rat, highlighting a role of the JAK2/STAT3 pathway in disease pathogenesis. Additionally, experimental trauma-hemorrhage results in a significant activation of the JAK2/STAT3 pathway in the liver and kidneys. Administration of baricitinib during resuscitation after major hemorrhage abolished the activation of JAK2/STAT3 as well as the activation of NF-κB and the NLRP3 inflammasome (liver and kidney); both of which are major drivers of local and systemic inflammation. Therefore, we propose that JAK inhibitors, such as baricitinib, may be repurposed to reduce the organ injury and inflammation caused by severe hemorrhage and resuscitation in patients with trauma.

## Abbreviations

ALT: alanine aminotransferase
AST: aspartate aminotransferase
CK: creatine kinase
DAMP: damage-associated molecular pattern
HS: hemorrhagic shock
JAK: Janus kinase
LDH: lactate dehydrogenase
MAP: mean arterial pressure
MODS: multiple organ dysfunction syndrome
STAT: signal transducer and activator of transcription

---

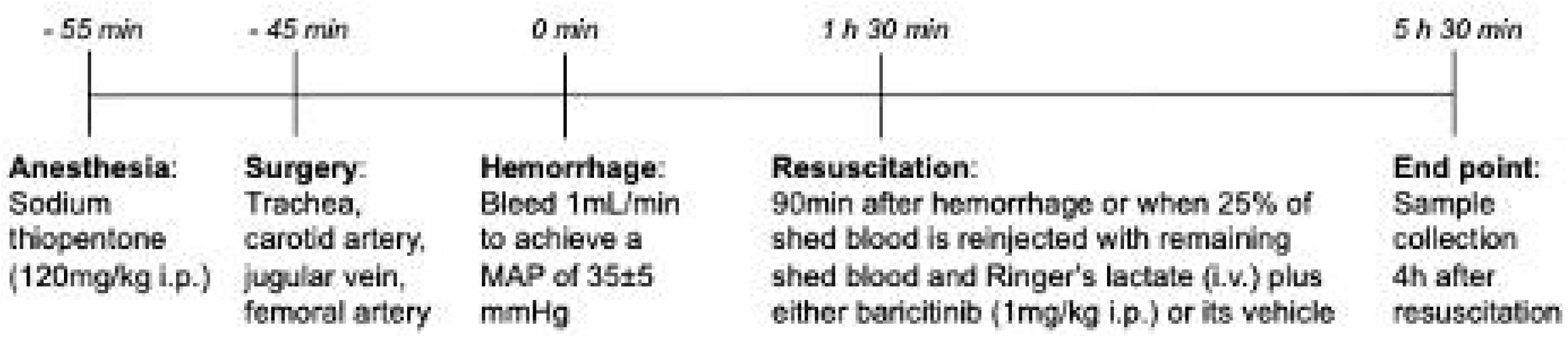

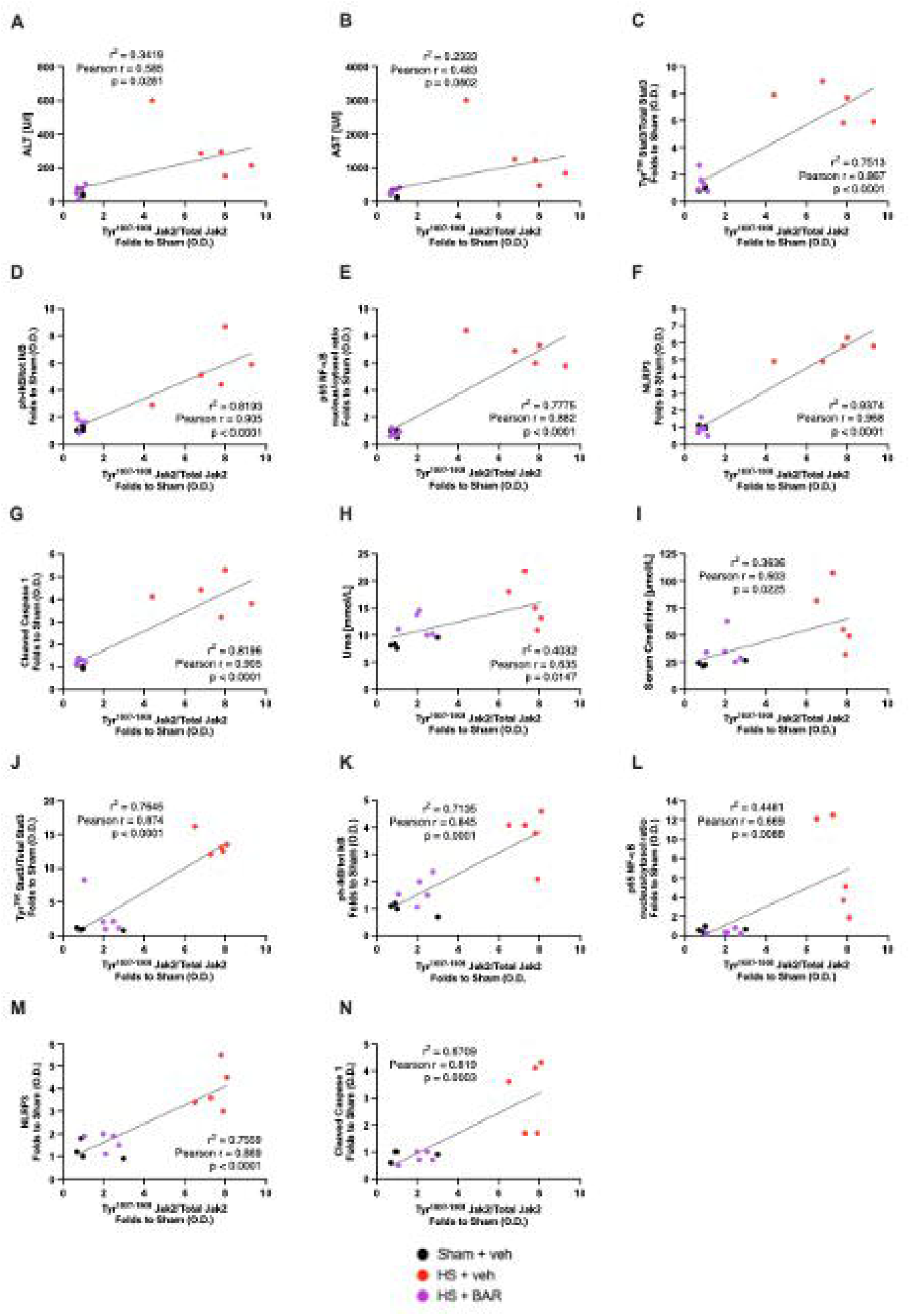

