## Supplemental for "Inhibition of the JAK/STAT pathway with baricitinib reduces the multiple organ dysfunction caused by hemorrhagic shock in rats"

### Supplemental Methods

#### Hemorrhagic Shock Model

Rats were anesthetized with sodium thiopentone (120 mg/kg i.p. initially and 10 mg/kg i.v. for maintenance as needed). Cannulation with polyethylene catheters (Smiths Medical International Ltd., Kent, UK) of the trachea for facilitation of spontaneous breathing (internal diameter [ID] 1.67 mm), left femoral artery for recording of the mean arterial pressure (MAP) (ID 0.40 mm), left carotid artery for blood withdrawal (ID 0.58 mm) and right jugular vein for fluid administration (ID 0.40 mm) was performed. To prevent tissue desiccation, swabs moistened with saline were placed over the surgery incision sites. Body temperature was monitored by a rectal probe thermometer and maintained at 37°C ± 0.3°C by means of a homoeothermic blanket system (Harvard Apparatus). Upon the completion of surgery, the MAP was allowed to stabilize for 15 min. Blood was then withdrawn (up to 1 mL/min into heparinized syringes containing 100 IU/mL heparin mixed with normal saline) through the cannula inserted in the carotid artery in order to achieve a fall in MAP to 35 ± 5 mmHg, which was recorded with a pressure transducer (attached to the femoral artery cannula, 844-31 Memscap, Durham, USA) and coupled to a PowerLab 8/30 data acquisition system (AD Instruments Pty Ltd., Castle Hill, Australia). Thereafter, MAP was maintained at 35 ± 5 mmHg for a period of 90 min either by further withdrawal of blood during the compensation phase or administration of the shed blood during the decompensation phase. At 90 min after initiation of hemorrhage (or when 25% of the shed blood had to be re-injected to sustain MAP at 35 ± 5 mmHg), resuscitation through the jugular vein was performed with the remaining shed blood (mixed with 100 IU/mL heparinized saline) over a period of 5 min plus a volume of Ringer's lactate identical to the volume of shed blood. Treatment or vehicle was administered intraperitoneally. One hour after resuscitation, an infusion of Ringer's lactate (1.5 mL/kg/h) was started as fluid replacement therapy, and it was maintained throughout the experiment for a total of 3 h. Under deep anesthesia, the heart was removed to terminate the experiment 4 h after resuscitation. Sham-operated rats were used as control and underwent identical surgical procedures, but without hemorrhage or resuscitation.

Rats were treated with either baricitinib (1 mg/kg) or its vehicle (5% DMSO + 95% Ringer's lactate) intraperitoneally as a bolus treatment immediately after resuscitation. This dose of baricitinib was based on preliminary optimal dose finding experiments.

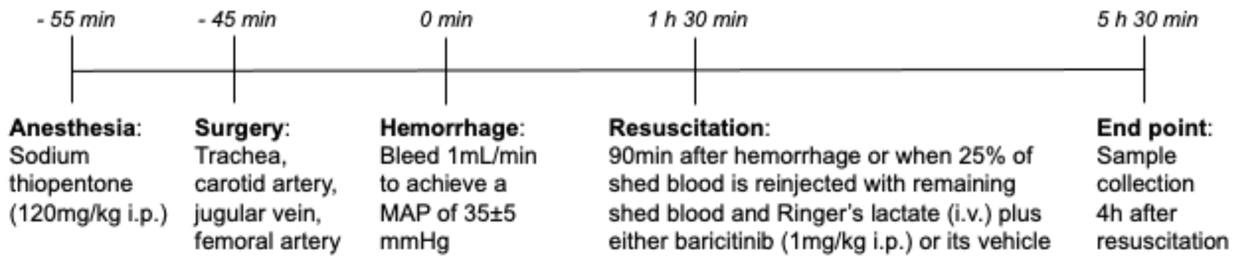


**Supplementary eFigure 1: Schematic representations of the acute HS model.** The experimental procedures at each stage of the acute HS model are shown.

**Sample Collection - Hemorrhagic Shock Model**

Rats remained anesthetized with sodium thiopentone (120 mg/kg i.p.) before sacrifice. Up to 5 mL of blood was taken from the carotid artery via the inserted cannula into a non-heparinized 5 mL syringe and immediately decanted into 1.1 mL serum gel tubes (Sarstedt, Germany). The blood was centrifuged (10,000 g for 5 min) to obtain the serum, which was subsequently stored at -80 °C until analysis. Organs (heart, lungs, liver, spleen and kidneys) were excised of which one section was snap frozen in liquid nitrogen and stored at -80 °C, and another section was placed in 10 % formalin for 24-48 h; followed by transfer to 70 % ethanol until further analysis. All organ injury/dysfunction parameters (urea, creatinine, alanine aminotransferase [ALT], aspartate aminotransferase [AST], creatine kinase [CK], amylase and lactate dehydrogenase [LDH]) in the serum were measured in a blinded fashion by a clinical pathology diagnostic laboratory (MRC Harwell Institute, Oxfordshire, UK).

**Western Blot Analysis**

Semi-quantitative immunoblot analysis were carried out in liver and kidney tissue samples as previously described[1,2]. Briefly, liver and kidney samples were homogenized in buffer and centrifuged (1320 g, 5 min, 4 °C). To obtain the cytosolic fraction, supernatants were centrifuged (16,125 g, 4 °C, 40 min). The pelleted nucleoli were resuspended in extraction buffer and centrifuged (16,125 g, 20 min, 4 °C). Protein content was determined on both nuclear and cytosolic extracts using bicinchoninic acid (BCA) protein assay (Thermo Fisher Scientific Inc, Rockford, IL). Proteins were separated by 8% sodium dodecyl sulfate polyacrylamide (SDS-PAGE) gel electrophoresis and electrotransferred to polyvinylidene difluoride (PVDF) membrane. After blocking (1 h in 10% dry milk solution), membranes were incubated with primary antibodies in 5% blocking solution overnight [1:1000 rabbit anti-NF-κB, 1:1000 mouse anti-$Ser^{32/36}$ IκBα, 1:1000 mouse anti-total IκBα, 1:1000 rabbit anti-$Tyr^{1007-1008}$ Jak2, 1:1000 rabbit anti-total Jak2, 1:1000 rabbit anti-$Tyr^{705}$ Stat3, 1:1000 mouse anti-total Stat3 (from Cell Signaling), 1:1000 rabbit anti-NLRP3 inflammasome (from Abcam), 1:1000 mouse anti-caspase 1 (p20) (from Adipogen)] followed by incubation with appropriate HRP-conjugated secondary antibodies. Proteins were detected with an ECL detection system and quantified by densitometry using analytic software (Quantity-One; Bio-Rad, Hercules, CA). Results were normalized with respect to densitometric values of tubulin for cytosolic proteins or histone H3 for nuclear proteins.

**CD68 immunohistochemical staining**

Endogenous peroxidase activity was blocked using 3% $H_2O_2$ (Carl Roth, Karlsruhe, Germany) for 20 min. A target retrieval solution (pH 6; Dako, Glostrup, Denmark) was used and antigen retrieval was performed for 10 min at 110 °C using a pressure cooker. Bovine serum albumin (BSA; Sigma Aldrich, St. Louis, MO, USA) in addition to avidin and biotin solution (Vector Laboratories, Burlingame, CA, USA) were used to block unspecific binding sites. Renal sections were incubated with anti-CD68 (1:500; abcam, Cambridge, United Kingdom) in 1% BSA in washing buffer overnight at 4 °C. Sections were further incubated with biotinylated anti-rabbit IgG (1:200; Vector Laboratories, Burlingame, CA, USA) and with VectaStain ABC kit (Vector Laboratories, Burlingame, CA, USA) according to the manufacturer's recommendations for 30 min. Substrate 3,3-diaminobenzidine (DAB; Vector Laboratories, Burlingame, CA, USA) was used and sections were counterstained with hemalaun (Carl Roth, Karlsruhe, Germany). Finally, renal sections were dehydrated and mounted for imaging. Tris(hydroxymethyl)aminomethan (TRIS) buffer (pH 7.6) containing 50 mM TRIS (Carl Roth, Karlsruhe, Germany), 300 mM sodium chloride (Carl Roth, Karlsruhe, Germany), 0.04% Tween® 20 (Sigma Aldrich, St. Louis, MO, USA) was used to wash renal sections during the staining process. CD68 staining was quantified by counting positive caskets of a grid (10 x 10) in 20 adjacent cortical areas per section (40x magnification). Images were taken using a KEYENCE BZ-X800 microscope and BZ-X800 viewer after performing white balance and auto exposure.

### Supplemental Information

#### Baricitinib

Baricitinib is a selective, reversible inhibitor of the JAK1 and JAK2 isoforms. As determined by isolated enzyme assays, baricitinib inhibits JAK1 and JAK2 almost to an equivalent degree; with $IC_{50}$ values of 5.9 nM and 5.7 nM, respectively. Baricitinib inhibits the JAK3 and TYK2 isoforms to a much lower extent; with $IC_{50}$ values of >400 nM and 53 nM, respectively. The rationale for developing a selective JAK1/2 inhibitor that does not inhibit JAK3 activity was to lower the immunosuppressive effects linked to pan-JAK inhibition[3,4].

Following the stimulation of cell surface receptors and subsequent activation of JAK isoforms, STAT proteins are phosphorylated and activated, leading to the activation of gene expression within the cell. Baricitinib modulates these signaling cascades through the inhibition of JAK1 and JAK2 which in turn decreases STAT protein phosphorylation and activation.

The half-life of baricitinib in beagle dogs is ~3.5 h and ~3-3.5 h in rats. In beagle dogs, no physiologically relevant changes were observed in blood pressure or heart rate parameters at any of the doses investigated[4].

Clinically, baricitinib is used for the treatment of rheumatoid arthritis and atopic dermatitis. The half-life of baricitinib in patients with rheumatoid arthritis is 12.5 h, whilst the half-life is 12.9 h in patients with atopic dermatitis. With regards to the patient factor differences (age, gender, race, ethnicity and body weight) following baricitinib treatment (humans), no clinically relevant effects on the pharmacokinetic properties of baricitinib were observed[3,4].

In November 2020, baricitinib in combination with remdesivir was granted an FDA Emergency Use Authorization for the treatment of suspected or laboratory confirmed COVID-19 in certain cohorts of hospitalized patients who require supplemental oxygen, invasive mechanical ventilation or extracorporeal membrane oxygenation[5].

### Supplemental Figures

#### JAK activation correlates with hepatic injury, renal dysfunction, STAT3, NF-κB and NLRP3 activation in an acute HS model

Firstly, to address the question whether the degree of activation of JAK correlates with changes in liver function, we correlated the degree of hepatic phosphorylation of JAK2 at $Tyr^{1007-1008}$ with serum ALT (Supplementary eFigure 2A) and serum AST (Supplementary eFigure 2B). We found a significant positive correlation between the degree of JAK activation and the increase in serum ALT (Supplementary eFigure 2A), suggesting that JAK activation drives or precedes the hepatic injury associated with HS. No significant correlation was observed between JAK activation and the increase in serum AST (Supplementary eFigure 2B), but AST is not a liver-specific enzyme. Secondly, the potential relationship between the degree of JAK activation and alterations in the activation of STAT and NF-κB were also addressed by correlating the degree of phosphorylation of JAK2 at $Tyr^{1007-1008}$ with the phosphorylation of STAT3 at $Tyr^{705}$ (Supplementary eFigures 2C [liver] and 2J [kidney]), phosphorylation of IκBα at $Ser^{32/36}$ (Supplementary eFigures 2D [liver] and 2K [kidney]) and the translocation of p65 (Supplementary eFigures 2E [liver] and 2L [kidney]). We found a highly significant positive correlation between the degree of JAK activation and STAT activation when measured as STAT3 phosphorylation (Supplementary eFigures 2C and 2J). We also found a highly significant positive correlation between the degree of JAK activation and NF-κB activation when measured as IκBα phosphorylation (Supplementary eFigures 2D and 2K) and p65 translocation (Supplementary eFigures 2E and 2L). Thirdly, whether the degree of JAK activation correlates with changes in the assembly and activation of the NLRP3 inflammasome was investigated by correlating the degree of phosphorylation of JAK2 at $Tyr^{1007-1008}$ with the expression of NLRP3 (Supplementary eFigures 2F [liver] and 2M [kidney]) and cleaved caspase 1 (Supplementary eFigures 2G [liver] and 2N [kidney]). We found a highly significant positive correlation between the degree of JAK activation and the NLRP3 inflammasome expression (Supplementary eFigures 2F and 2M) and activation of caspase 1 (Supplementary eFigures 2G and 2N). Lastly, we correlated the degree of renal phosphorylation of JAK2 at $Tyr^{1007-1008}$ with serum urea (Supplementary eFigure 2H) and serum creatinine (Supplementary eFigure 2I) to assess whether the degree of activation of JAK correlates with changes in kidney function. We found a significant positive correlation between the degree of JAK activation and the increases in serum urea (Supplementary eFigure 2H) and creatinine (Supplementary eFigure 2I), suggesting that JAK activation drives or precedes the renal dysfunction associated with HS.

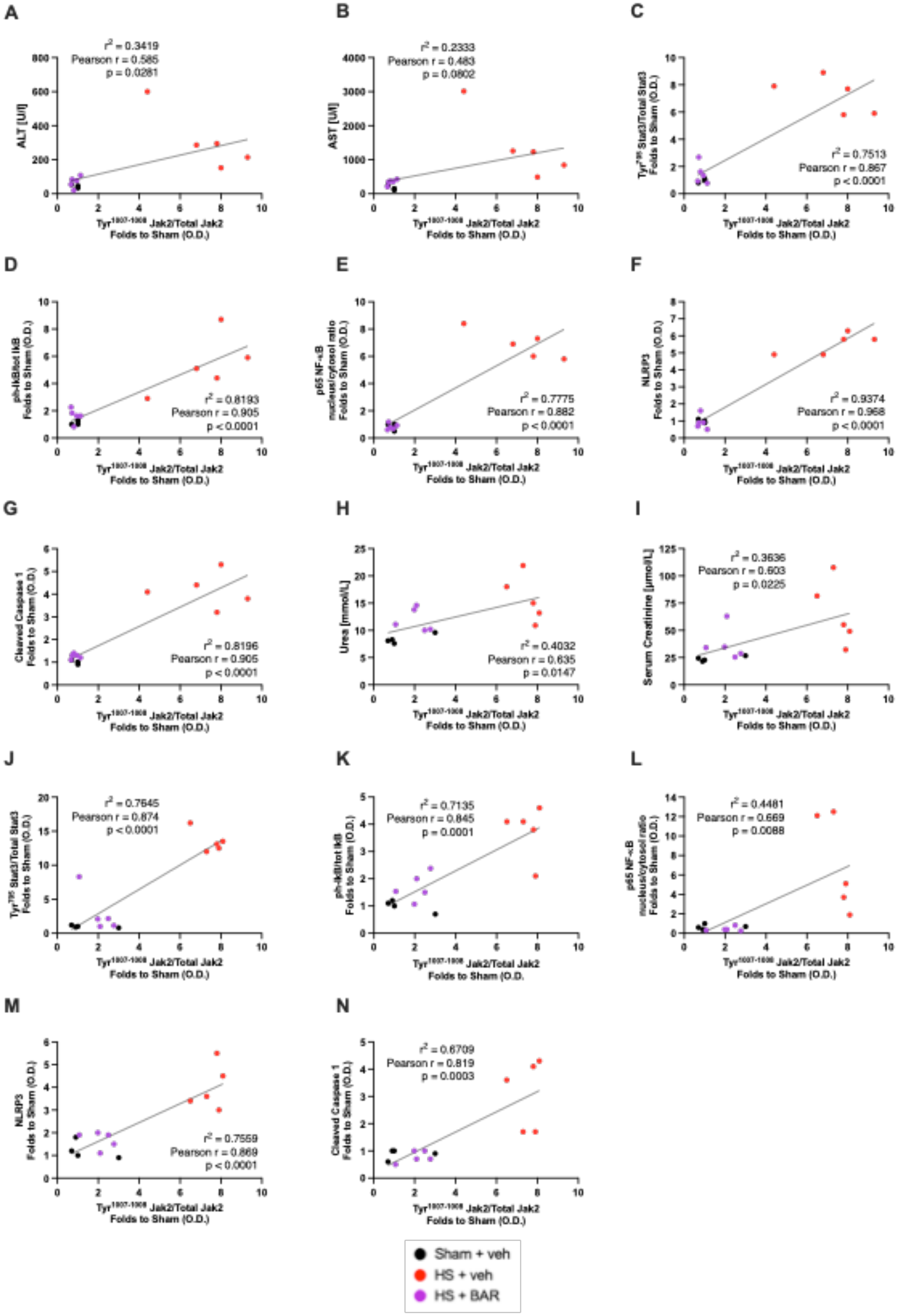
A
$r^2 = 0.3419$
Pearson r = 0.585
p = 0.0261
ALT [U/l]
B
$r^2 = 0.2333$
Pearson r = 0.483
p = 0.0802
AST [U/l]
C
$r^2 = 0.7513$
Pearson r = 0.867
p < 0.0001
Tyr705 Stat3/Total Stat3 Folds to Sham (O.D.)
D
$r^2 = 0.8193$
Pearson r = 0.905
p < 0.0001
ph-IkB/tot IkB Folds to Sham (O.D.)
E
$r^2 = 0.7775$
Pearson r = 0.882
p < 0.0001
p65 NF-κB nucleus/cytosol ratio Folds to Sham (O.D.)
F
$r^2 = 0.9374$
Pearson r = 0.968
p < 0.0001
NLRP3 Folds to Sham (O.D.)
G
$r^2 = 0.8196$
Pearson r = 0.905
p < 0.0001
Cleaved Caspase 1 Folds to Sham (O.D.)
H
$r^2 = 0.4032$
Pearson r = 0.635
p = 0.0147
Urea [mmol/L]
I
$r^2 = 0.3636$
Pearson r = 0.603
p = 0.0225
Serum Creatinine [µmol/L]
J
$r^2 = 0.7645$
Pearson r = 0.874
p < 0.0001
Tyr705 Stat3/Total Stat3 Folds to Sham (O.D.)
K
$r^2 = 0.7135$
Pearson r = 0.845
p = 0.0001
ph-IkB/tot IkB Folds to Sham (O.D.)
L
$r^2 = 0.4481$
Pearson r = 0.669
p = 0.0088
p65 NF-κB nucleus/cytosol ratio Folds to Sham (O.D.)
M
$r^2 = 0.7559$
Pearson r = 0.869
p < 0.0001
NLRP3 Folds to Sham (O.D.)
N
$r^2 = 0.6709$
Pearson r = 0.819
p = 0.0003
Cleaved Caspase 1 Folds to Sham (O.D.)
Tyr1007-1008 Jak2/Total Jak2 Folds to Sham (O.D.)
Sham + veh
HS + veh
HS + BAR

**Supplementary eFigure2: JAK activation correlates with hepatic injury, renal dysfunction, STAT3, NF-κB and NLRP3 activation in an acute HS model.** Linear regression analysis of (**A**) hepatic phosphorylation of JAK2 at $Tyr^{1007\text{-}1008}$ vs. serum ALT, (**B**) hepatic phosphorylation of JAK2 at $Tyr^{1007\text{-}1008}$ vs. serum AST, (**C**) hepatic phosphorylation of JAK2 at $Tyr^{1007\text{-}1008}$ vs. phosphorylation of STAT3 at $Tyr^{705}$, (**D**) hepatic phosphorylation of JAK at $Tyr^{1007\text{-}1008}$ vs. phosphorylation of IκBα at $Ser^{32/36}$, (**E**) hepatic phosphorylation of JAK at $Tyr^{1007\text{-}1008}$ vs. translocation of p65, (**F**) hepatic phosphorylation of JAK at $Tyr^{1007\text{-}1008}$ vs. expression of NLRP3, (**G**) hepatic phosphorylation of JAK at $Tyr^{1007\text{-}1008}$ vs. expression of the cleaved form of caspase 1, (**H**) renal phosphorylation of JAK at $Tyr^{1007\text{-}1008}$ vs. serum urea, (**I**) renal phosphorylation of JAK at $Tyr^{1007\text{-}1008}$ vs. serum creatinine, (**J**) renal phosphorylation of JAK2 at $Tyr^{1007\text{-}1008}$ vs. phosphorylation of STAT3 at $Tyr^{705}$, (**K**) renal phosphorylation of JAK at $Tyr^{1007\text{-}1008}$ vs. phosphorylation of IκBα at $Ser^{32/36}$, (**L**) renal phosphorylation of JAK at $Tyr^{1007\text{-}1008}$ vs. translocation of p65, (**M**) renal phosphorylation of JAK at $Tyr^{1007\text{-}1008}$ vs. expression of NLRP3 and (**N**) renal phosphorylation of JAK at $Tyr^{1007\text{-}1008}$ vs. expression of the cleaved form of caspase 1. Data are expressed as raw individual values of 4-5 animals per group. Statistical analysis was performed using simple linear regression to calculate the $r^2$ value, Pearson correlation coefficient test to calculate the r value and a two-tailed t-test to calculate the p-value. *$p<0.05$ denoted statistical significance.

### Supplemental Discussion

#### Macrophage recruitment

Following the recruitment of neutrophils and release of pro-inflammatory mediators, monocytes are attracted to the site of injury by chemotaxis and differentiate into macrophages. Activated macrophages are phenotypically classified as either M1 pro-inflammatory or M2 anti-inflammatory macrophages[6]. From the resulting exposure to pro-inflammatory cytokines such as TNF-α and IFN-γ and cellular debris from apoptotic or necrotic cells, macrophages tend to be polarized more towards the classically activated M1 phenotype to stimulate an inflammatory response within the injured tissue[7–9]. M1 macrophages produce IL-1, IL-6, TNF-α and ROS which further recruit leukocytes to injured tissues and increases apoptosis, both of which contribute to organ inflammation. The observed improvement in histomorphological changes may in part be due to the reduced neutrophil and macrophage invasion and thus, lower level of excessive inflammation.

#### Limitations of the study

Although baricitinib demonstrated some striking, beneficial effects in the rat model of HS, there are limitations which should be taken into consideration. An acute model of HS was used in this study, which results in systemic inflammation and MODS within a few hours following resuscitation. At 4 h post resuscitation, there is organ injury and dysfunction and significant activation of JAK/STAT, NF-κB and NLRP3. Despite the effectiveness of baricitinib in this acute setting, we cannot conclude that the same protective effects will be observed in animal models with a longer follow-up period. Moreover, survival studies are needed to confirm that the observed early reduction in MODS does, indeed, translate to improved outcome and ultimately a decrease in mortality; all of which would strengthen the efficacy data of baricitinib in this study. Organ injury and dysfunction were used as surrogate markers for mortality in our study (as the determination of mortality is not allowed by our ethics and Home Office license). Thus, caution should be taken when interpreting our pre-clinical results and extrapolating them to the clinical scenario. In this study, gender differences were not investigated, as only male rats were used to prevent any sex-dependent confounding effects (variations in female reproductive hormones and X chromosome) and to represent the population usually most affected by trauma (young males). Of note, the majority of trauma patients in our recent TOP-ART clinical trial were male and aged under 30, so the use of male rats is clinically appropriate[10]. Further studies in larger animals (e.g. pigs) and/or higher species may be useful to increase understanding of both the mechanism of action (e.g. blood gas analysis and microcirculatory effects) and verify efficacy of baricitinib in HS. We show that treatment with baricitinib upon resuscitation had no significant effect on arterial blood pressure, suggesting that the beneficial influence of baricitinib in HS may be independent of effects on blood pressure and, hence, microvascular perfusion of the tissues/organs. This could potentially limit the effectiveness of baricitinib treatment in HS, as alternative therapies may be required to counteract the vascular decompensation associated with HS. Nonetheless, clinical studies with large cohorts of trauma patients are required to robustly examine the relationship between JAK activity and clinical outcomes in humans.
